# Infant Brains Tick at 4Hz – Resonance Properties of the Developing Visual System

**DOI:** 10.1101/2025.09.11.675068

**Authors:** Marlena Baldauf, Ole Jensen, Moritz Köster

**Author notes:** Correspondence: Marlena Baldauf Moritz Köster.

## Abstract

Neural rhythms of the infant brain are not well understood. Testing the rhythmic properties of the adult visual system with periodic or broadband visual stimulation elicited neural resonance phenomena at ∼10Hz alpha rhythm. To investigate the rhythmic properties of the infant brain upon visual stimulation, eight-month-olds (*N* = 42) were presented with visual stimuli flickering at discrete frequencies (2, 4, 6, 8, 10, 15, 20, and 30Hz) and broadband (i.e., aperiodic) stimulation, while recording a high-density electroencephalogram (EEG). Infants’ visual system entrained to the harmonics of the periodic stimulation frequencies (first to third). Crucially, a 4Hz rhythm emerged independent of stimulation frequency and the impulse response function (IRF) of the broadband sequence revealed a perceptual echo, reverberating visual information at 4Hz. This echo lasted for about 1 second, extended into frontal sensors, and selectively resonated the 4Hz component of the input signal. In a complementary adult sample, we confirm an alpha response upon periodic and broadband stimulation. To conclude, perturbing the infant visual system elicited a neural response and resonant activity at the 4Hz theta rhythm, which contrasts with the 10Hz alpha rhythm found in the adult visual system. Neural processing dynamics are thus essential to understand early brain development in full.

**Significance Statement:** This study tested resonance properties of the infant visual system upon perturbation. For the visual stimulation at discrete frequencies between 2 and 30Hz, the infant brain responded at 4Hz independent of the stimulation frequency. Following broadband stimulation, a perceptual echo at 4Hz emerged – resonating the 4Hz component of visual information over several cycles. These findings reveal that the 4Hz theta rhythm is an inherent processing dynamic of the infant visual system, contrasting with the 10Hz rhythm found in adults. The processing dynamics of the human visual system undergo fundamental changes from infancy into adulthood.

---

Human brain dynamics are characterized by neural rhythms, which coordinate information processing in distributed neural networks (Buzsáki, 2002; Engel et al., 2001; Fell & Axmacher, 2011; Fries, 2015; Hanslmayr et al., 2016; Palva & Palva, 2007; Varela et al., 2001). In the adult brain, neural processes in the 3-8Hz theta and the 8-14Hz alpha range have been studied in depth (e.g. Hanslmayr et al., 2016; Jensen & Mazaheri, 2010; Klimesch, 1997; Köster & Gruber, 2022). However, the emergence of neural rhythms in the infant brain and their trajectory towards the rhythmic processing dynamics found in adults remain unclear.

The alpha rhythm is the most dominant frequency in the adult brain (H. Berger, 1929; Klimesch, 1999) and is associated with attentional processes (Klimesch, 1999; Köster & Gruber, 2022). It is proposed to reflect the gating of information in perceptual networks, inhibiting task-irrelevant information when increased (i.e., pulsed inhibition) and promoting neural processing in perceptual and task-relevant cortical networks when reduced (Jensen & Mazaheri, 2010). Theta activity is associated with mnemonic functions (Axmacher et al., 2008; Fell et al., 2003; Friese et al., 2013; Köster et al., 2018), specifically with the ordering and maintenance of information in working memory (Jensen & Tesche, 2002; Lisman & Idiart, 1995) and facilitating long-term potentiation in medio-temporal networks (Buzsáki, 2002; Köster, 2024; Lisman & Idiart, 1995; Lisman & Jensen, 2013).

Neural rhythms have been intensely studied with respect to the encoding of visual information. But how do these neural dynamics emerge in the infant brain? There is initial evidence that the theta rhythm is present and may likewise serve a mnemonic function in infants (Angelini et al., 2023; Begus et al., 2015; Köster et al., 2021; Orekhova, 1999). Regarding alpha, a somewhat lower 6-9 Hz rhythm was modulated by attentional processes in nine-month-olds, akin to the suppression of the alpha rhythm in adults (Hoehl et al., 2014). However, a systematic investigation of infant neural processing dynamics upon visual stimulation is missing.

A key approach to investigate the dynamic properties of the brain is to perturbate the visual system by rhythmic stimulation (RVS), periodically at discrete frequencies, or aperiodically using broadband luminance sequences (Adrian & Matthews, 1934; Clouter et al., 2017; Herrmann, 2001;

Köster et al., 2019; Norcia et al., 2015; VanRullen & Macdonald, 2012). In the visual domain, RVS involved the presentation of stimuli (or strobe light) at different frequencies and the measurement of rhythmic brain responses in the magneto- and electroencephalogram (M/EEG; Adrian & Matthews, 1934; Köster et al., 2023; Norcia et al., 2015). This commonly elicits brain responses at the stimulation frequency (i.e. first harmonic) as well as its multiples and fractions (i.e. higher harmonics and subharmonics, respectively). In addition, the responses in some frequencies might be amplified, reflecting resonant properties of the neuronal system at that frequency (Buzsáki & Wang, 2012). Such resonant properties have been taken as markers for the endogenous dynamical properties of the visual system (Herrmann, 2001; VanRullen & Macdonald, 2012).

In the adult visual system, resonance phenomena have been identified mostly around 10Hz. In a seminal study, Hermann (2001) used strobe lights at frequencies ranging from 1 to 100Hz in 1Hz steps. Besides brain responses at harmonic frequencies, a consistent 10Hz alpha response emerged, independent of the stimulation frequency. Notably, stimulation near 10Hz elicited a resonance phenomenon, with the visual cortex responding more strongly to flicker at this frequency than to adjacent frequencies. Van Rullen and Macdonald (2012) used a broadband (i.e., aperiodic) visual stimulation. That is, they presented sequences of random luminance values tailored to have a flat frequency spectrum. This elicited a 10Hz impulse response function (IRF) in the corresponding EEG signal, suggesting that the broadband luminance sequence was repeated in the EEG signal, lasting for around one second, which they coined a *perceptual echo*. Critically, the 10Hz signal disappeared after removing the 10Hz component form the broadband signal, which indicates a frequency-specific resonance of the 10Hz component of visual inputs.

Here, we extend the RVS approach to uncover dynamic properties of the visual system in the seventh to ninth postnatal month. Specifically, we presented *N* = 42 8-month-olds with images of child-friendly cartoon monsters flickered periodically at discrete frequencies (i.e., appearing and disappearing) at 2, 4, 6, 8, 10, 15, 20 or 30Hz or aperiodically at broadband sequences (luminance varying frame-by-frame with an overall flat frequency spectrum), while measuring a 64-channel active EEG (see Figure 1). Images were presented for 2s in combination with randomized ‘monster sounds’. Given the prominence of the theta rhythm in infancy (Cellier et al., 2021; Köster et al., 2021; Xie et al., 2022), we expected the presence of a 4Hz theta response and resonant activity in the infant visual system upon periodic stimulation and broadband visual input, respectively.

**Figure 1.**
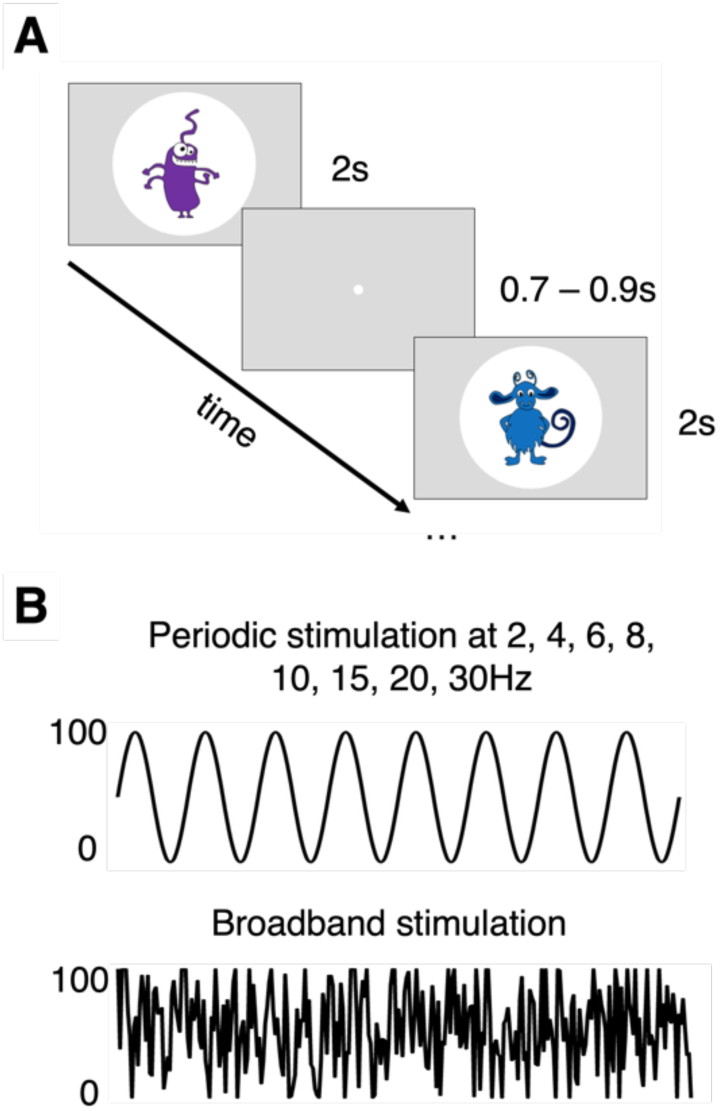
Example trial sequence and rhythmic visual stimulation. (A) In each trial, infants saw a cartoon monster inside a white circle for 2 s (ISI: 0.7 to 0.9s). (B) Stimuli were presented periodically at discrete eight frequencies (2, 4, 6, 8, 10, 15, 20, 30Hz) and in broadband sequences.

## Results

### Periodic visual stimulation revealed an endogenous 4Hz rhythm in the infant brain

We examined frequency spectra in response to periodic stimulation using Fast Fourier Transformation. The neural response to periodic stimulation is depicted in Figure 2 (see also Table 1). We obtained significant responses at all first harmonics, all *p* < .01 (SNR of the power at first harmonic frequency compared to SNR = 1), except for 2Hz (*p* = .999), due to power increase towards lower frequencies (i.e., 1/f). Furthermore, a significant second harmonic (2 times the first harmonic frequency) emerged for all tested frequencies, all *p* < .001 (out of range: 30Hz). A significant third harmonic (3 times the first harmonic frequency) was present for most stimulation frequencies (4, 6, 8 and 10Hz), all *p* < .001, apart from 2Hz, *p* = .081 (out of range: 15, 20 and 30Hz). Topographically, the harmonic responses peaked at occipital sites (Figure 2B).

**Figure 2.**
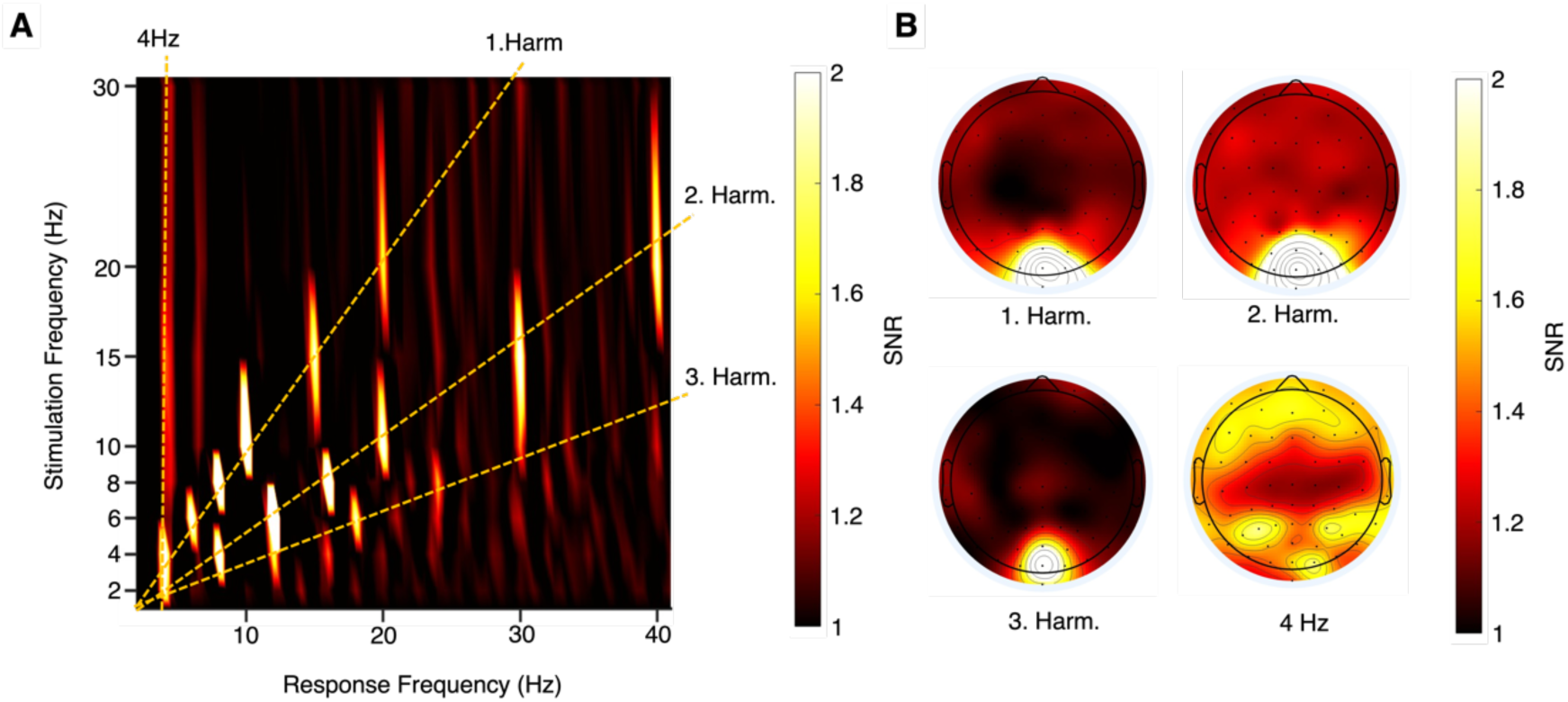
Visual system response to periodic stimulation at 2, 4, 6, 8, 10, 15, 20, 30Hz. (A) The neural response spectra as a function of stimulation frequencies at parietooccipital electrodes. The signal-to-noise ratio (SNR) was above chance (SNR = 1) for the first, second and third harmonic responses, and the 4Hz rhythm for most stimulation frequencies (see Table 1 for statistics). (B) Topographic maps display averaged responses at first, second and third harmonic, and at 4Hz across all stimulation frequencies.

**Table 1.** Visual system response to periodic stimulation at 2, 4, 6, 8, 10, 15, 20, 30Hz. Mean SNRs and standard deviations are presented for the first, second and third harmonics harmonic, and the ∼4Hz response. For the 4Hz response, the maximum frequency (i.e., 4 or 4.5Hz) is reported. Statistics are based on one-sample t-tests against baseline (SNR = 1). * significant at p < .05, ** significant at p < .01, ***significant at p < .001.

| Stimulation<br>frequency | Signal to Noise Ratio |  |  |  |
| --- | --- | --- | --- | --- |
|  | 1. Harmonic | 2. Harmonic | 3. Harmonic | ~4Hz |
| 2Hz | 0.646(0.269) | 2.178(1.35)*** | 1.1(0.452) | 2.178(1.35)<sup>4</sup>*** |
| 4Hz | 2.542(2.258)*** | 2.906(1.982)*** | 1.838(0.96)*** | 2.542(2.258)<sup>4</sup>*** |
| 6Hz | 2.464(1.746)*** | 7.162(7.290)*** | 2.038(1.371)*** | 1.138(0.281)<sup>4</sup>*** |
| 8Hz | 3.829(3.392)*** | 3.524(2.004)*** | 1.568(0.863)*** | 1.302(0.561)<sup>4.5</sup>*** |
| 10Hz | 3.811(2.303)*** | 2.404(1.242)*** | 1.489(0.474)*** | 1.311(0.495)<sup>4.5</sup>*** |
| 15Hz | 2.091(1.061)*** | 2.311(1.282)*** | n/a | 1.434(0.688)<sup>4</sup>*** |
| 20Hz | 1.763(0.91)*** | 2.097(1.299)*** | n/a | 1.212(0.476)<sup>4.5</sup>** |
| 30Hz | 1.201(0.472)** | n/a | n/a | 1.241(0.389)<sup>4.5</sup>*** |

In addition, we found a highly significant 4Hz or 4.5Hz response (all *p* <. 001; see Table 1), independent of stimulation frequency. This 4Hz response was similarly present in the no-flicker condition, *t*(41) = 3.287, *p* = .001, but not in the omission condition, *t*(40) = 1.460, *p* = .08 (both presented as control conditions, see methods for details). The topography of this 4Hz response (Figure 2B) showed a widespread distribution at parietooccipital and frontal recording sites. The 4Hz response was particularly pronounced for frontal sites (see Figure S1), where it showed a higher response than first harmonic responses, *t*(41) = -3.599, *p* < .001, and also compared to the frontal theta activity for the no-flicker and omission condition *t*(40) = -3.606, *p* < .001. No such response emerged in any frequency within the alpha range (8-14Hz; except for harmonics and their adjacent frequency steps).

Spectral representations and topographical distributions for all stimulation frequencies and the responses to the no-flicker and omission condition are provided as supplemental materials (Figure S2 and S3).

### Broadband visual stimulation elicits a 4Hz resonant response in the infant brain

The Impulse Response Function (IRF) of the broadband sequences and the corresponding EEG signals was computed as their cross-correlation (see Methods for details). For comparison, we computed a surrogate function as the cross-correlation of the EEG signal with shuffled broadband sequences of other infants. The IRF of the broadband stimulation showed a rhythmic 4Hz pattern between 0 and 1s after stimulus onset (Figure 3B; red). This 4Hz response was also visible in the corresponding time-frequency representation of power (TFR) and in contrast to the surrogate, shown in blue in Figure 3B. Relative to the surrogate, the IRF showed increased 4 Hz power in the 0 to 1 s window, *t*(41) = -12.698, *p* < .001 (Figure 3C). The topography of the 4Hz IRF response was widespread with maximal power at parietooccipital and frontocentral sites (Figure 3D).

**Figure 3.**
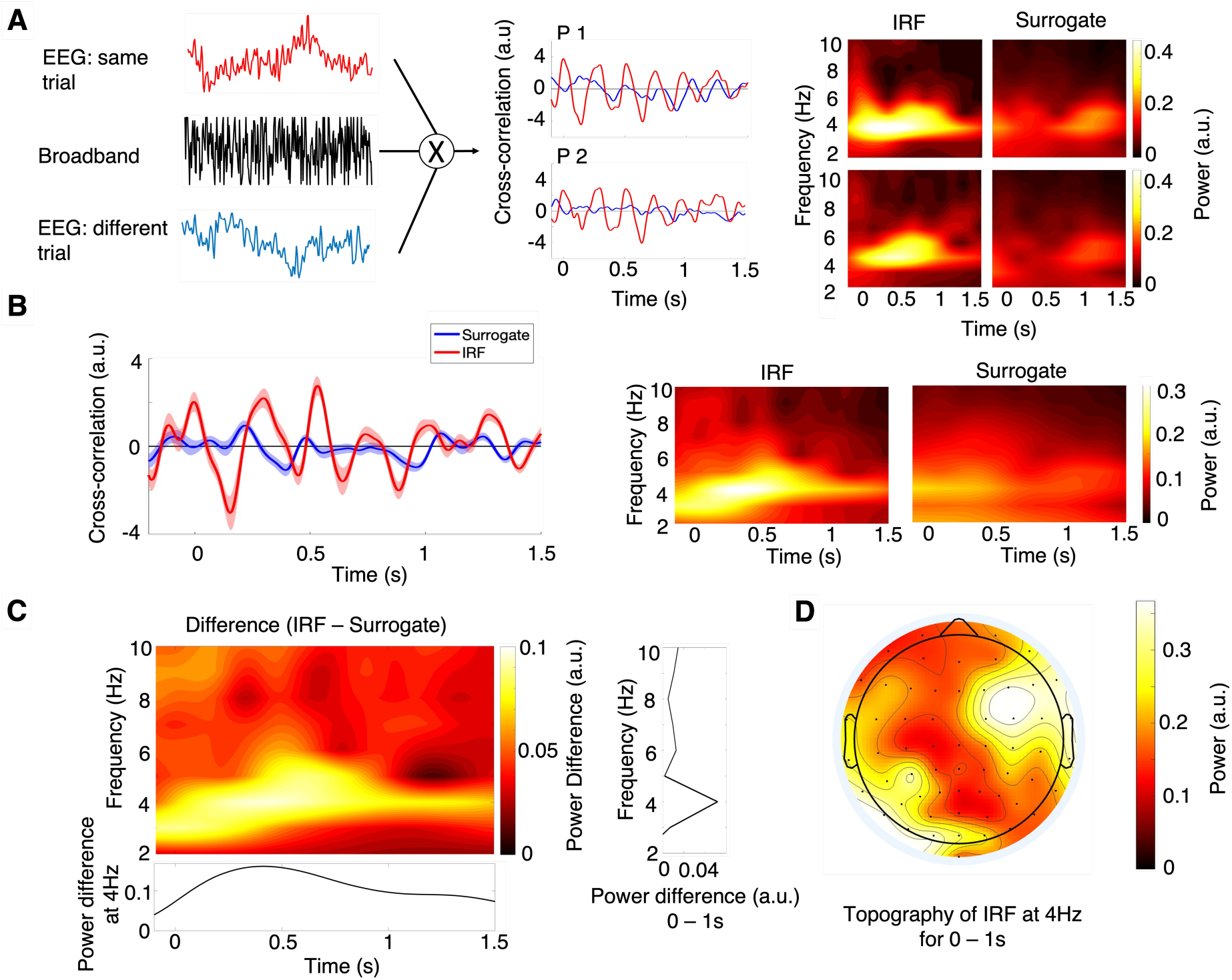
Visual system response to broadband visual stimulation. (A) The IRF was estimated by cross-correlating the broadband sequences and the EEG signal of the same trials. The surrogate was calculated as the cross-correlation with random trials (blue). The IRF, surrogate and corresponding spectral representations are shown for two representative participants (P). (B) IRF and surrogate together with their spectral representation across all infants. (C) The spectral difference between the IRF and the surrogate and (D) the topography of the IRF at 4Hz.

### The infant brain selectively reverberates the 4Hz component of the broadband signal

The IRF of the band-stop filtered broadband sequence show a marked reduction in 4 Hz power, when compared to the IRF of the original signal, as confirmed by a paired samples t-test, *t*(41) = 17.826, *p* < .001 (Figure 4).

**Figure 4.**
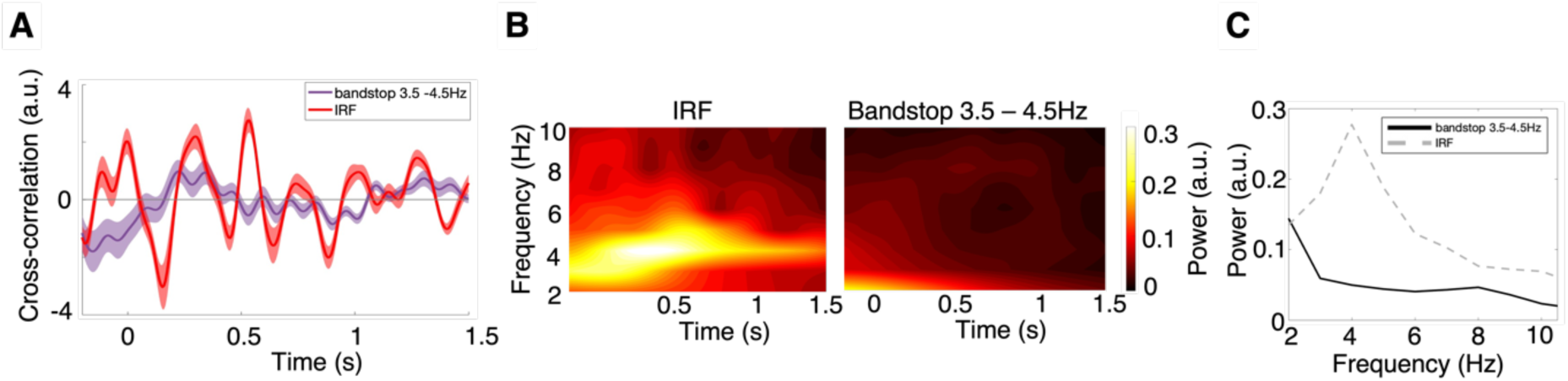
Response to broadband visual stimulation after removing the 3.5 to 4.5Hz component of the broadband sequence. (A) IRF of the band-stop filtered stimulus sequence (purple) compared to the IRF of the full-spectrum stimulus (red). (B) The corresponding TFRs show reduced 4 Hz power after filtering. (C) Power spectra (2–10 Hz) of the 0 to 1 s time window for the band-stop filtered and full spectrum IRFs.

### In adults, visual stimulation revealed an alpha response

For direct comparison within the same paradigm, we also tested a small adult sample (*N* = 7) and observed significant first harmonic responses at all stimulation frequencies, *p* < .05, except 2Hz, *p* = .083 (Figure 5A). Likewise, the second harmonic emerged for all tested stimulation frequencies, *p* < .05 (out of range: 30Hz) and significant third harmonic for all tested stimulation frequencies, *p* < .05, with marginal significance for 4Hz, at *p* = .068 (out of range: 15, 20, 30Hz). Figure 5B depicts the topographical distribution of harmonic responses peaking at occipital sites.

**Figure 5.**
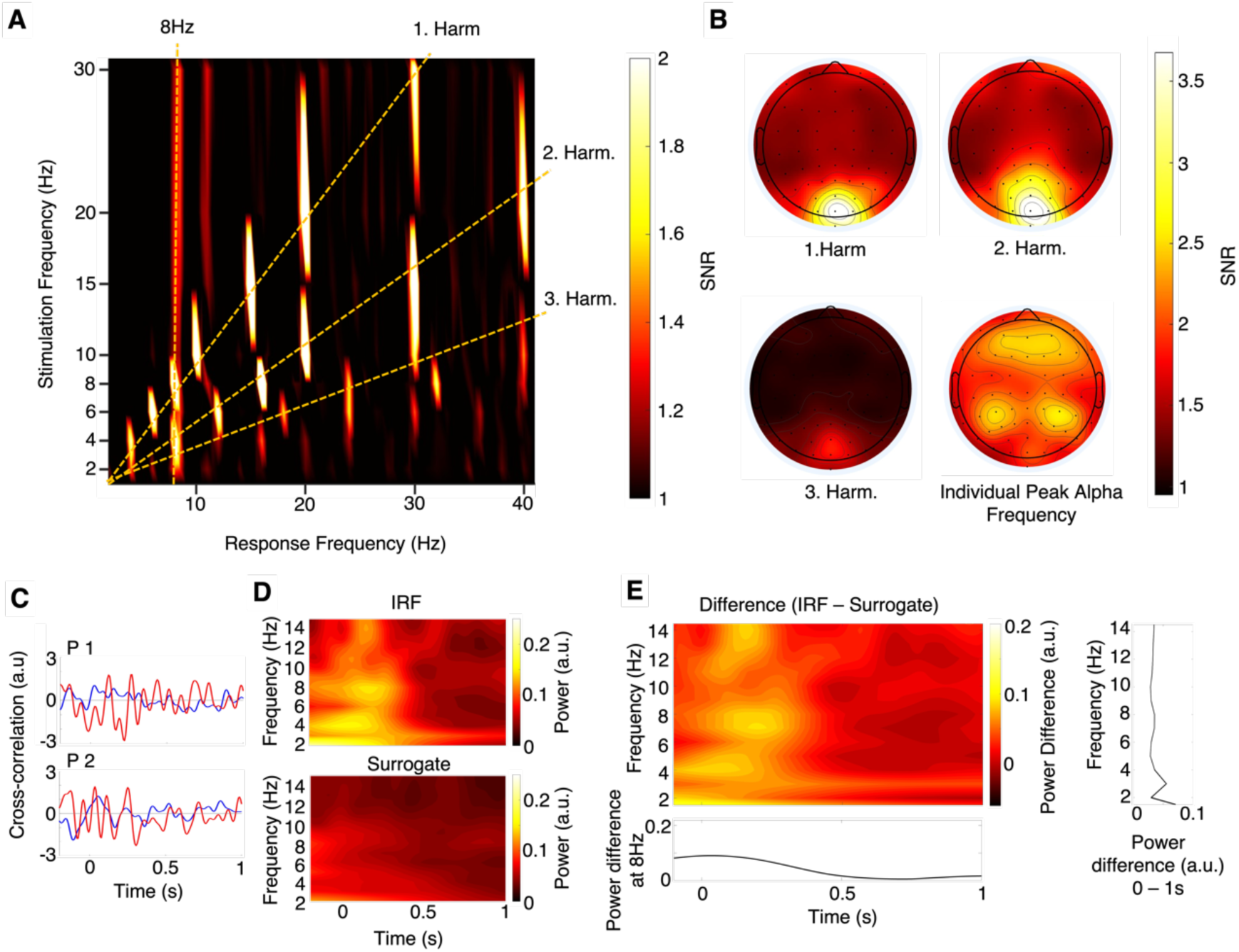
Visual system response to periodic stimulation at 2, 4, 6, 8, 10, 15, 20, 30Hz and to broadband stimulation in N = 7 adults. (A) The neural response spectra as a function of stimulation frequencies at parietooccipital electrodes. The SNR was above chance (SNR = 1) for first, second and third harmonic responses, and for the individual peak alpha frequency for most stimulation frequencies. (B) Topographic maps display averaged responses at first, second and third harmonic, and at the individual peak alpha frequency across all stimulation frequencies. (C) Response to broadband stimulation for two representative participants - IRF (red) and surrogate (blue) (D). TFR of IRF (top) and the surrogate function (bottom) across all adult subjects and (E) their difference.

We observed a significant increase in the individual peak alpha frequency (8-14Hz) following stimulation at 2, 4, 6, 8, 20Hz, all *p* < .05 (one-tailed tests; due to the predicted increase) and a trend for 10, 15, and 30Hz, all *p* < .071, suggesting an endogenous rhythm at the alpha frequency. We found an average peak alpha frequency of 8.57Hz, as indicated by the vertical line in Figure 5A. The topographical distribution of the alpha response was widespread across all recoding sites (Figure 5B).

Figure 5C shows the response to broadband stimulation (IRF) for two representative subjects (red) alongside their surrogate functions (blue). The TFRs of IRF and surrogate across all *N* = 7 subjects are illustrated in Figure 5D. The IRF TFR exhibits oscillatory responses at multiple frequencies that are absent in the surrogate TFR (see also Figure 5E for the IRF – surrogate difference). A paired samples t-test revealed a significant difference in peak alpha power between the IRF and surrogate, *t* (6) = 3.5197, *p* = .006, with greater power in the IRF. In contrast, no significant differences between the IRF and surrogate were observed at 4Hz, *t*(6) = -2.015, *p* = .091, or at adjacent frequencies (3.5Hz and 4.5Hz), both *p* > .130.

## Discussion

Rhythmic stimulation of the infant visual system revealed a pronounced 4Hz theta response. This was evident, first, by an endogenous 4Hz theta rhythm that emerged during periodic stimulation (2, 4, 6, 8, 10, 15, 20, 30Hz) and, second, a resonance phenomenon at 4Hz following broadband stimulation. This suggests that the 4Hz theta rhythm constitutes an inherent processing dynamic of the infant visual system, which stands in stark contrast to the resonance phenomena in the alpha range observed in the adult visual system.

For periodic stimulation, a consistent 4Hz response emerged independent of stimulation frequency. Critically, this was also the case for frequencies without 4Hz harmonics, namely the 6, 10, 15 and 30Hz stimulation. Thus, the 4Hz rhythm in the infant brain is robust against external rhythmic pacemakers and constitutes an endogenous oscillation. Topographically, the 4Hz response was widely distributed, being accentuated at parietooccipital and frontocentral sites. As expected, we also observed increased power at the first, second and third harmonics, most prominently at occipital electrodes.

Broadband stimulation revealed an IRF at 4Hz that lasted for at least 1 second, suggesting that the infant visual system *reverberates* (i.e. echoes) at 4Hz frequency. This echo was found at occipital sensors and in frontocentral recording sites, which are prominent for their strong theta response in adults (Cavanagh & Frank, 2014; Friese et al., 2013). Crucially, when removing the 4Hz component from the broadband signal, this echo disappeared, indexing that the infant brain selectively resonated the 4Hz component of the broadband input.

Taken together, these findings indicate that the theta rhythm constitutes an inherent rhythmic property of the infant visual system, which stands in contrast with the resonance phenomena at the alpha rhythm found in adults in former studies (Herrmann, 2001; VanRullen & Macdonald, 2012) as well as our adult sample, replicating former findings using the present design. The observed shift from the theta to alpha rhythm from infancy to adulthood corroborates findings from the resting-state literature (Cellier et al., 2021; Wilkinson et al., 2024). There are two accounts of the shift in brain dynamics from infancy into adulthood – a *mnemonic* and an *inhibition* account.

According to a mnemonic account, the theta rhythm in infants is functionally equivalent to the adult theta rhythm (Köster et al., 2021; Köster & Gruber, 2022). Crucial to this argument is the observation that, first, theta activity is associated with the formation of novel representations across development: in infants (Begus et al., 2015; Xie et al., 2022), children (Köster et al., 2017), and adults (Friese et al., 2013; Köster et al., 2018), and second, theta power increases in response to novel information infants (A. Berger & Posner, 2023; Köster et al., 2019, 2021) and adults alike (Cavanagh &

Frank, 2014). The developmental shift in the dominant frequency from theta to alpha may thus reflect a developmental change in the operating mode of the brain from infancy to adulthood: In adults, decreased alpha activity is suggested to reflect the gating of perceptual information in established cortical networks (Klimesch, 2012; Köster & Gruber, 2022). In contrast, infants cannot yet rely on such mature cortical networks, because these are still ‘in the making’. Instead, information may be processed in broader networks, including frontal and medio-temporal sites, supporting the formation and refinement of cortical networks that allow for fast and efficient processing in older children and adults (Buzsáki, 2002; Köster, 2024; Lisman & Jensen, 2013) . In support of this notion, there is recent evidence that infants as young as three months rely on the hippocampus during statistical and perceptual learning (Ellis et al., 2021; Yates et al., 2025). Thus, the prominent theta dynamics in the infant brain may reflect the constant sampling and updating processes that guide early concept formation.

The inhibition account of the theta rhythm and perceptual echo in infants proposes that oscillatory dynamics in the theta band serve a similar role to the alpha rhythm in adults. According to this view, theta oscillations act as a mechanism for pulsed inhibition, helping regulate the allocation of neural resources during sensory processing—for example, modulated by spatial attention. The developmental increase in oscillatory frequency is consistent with the observed decrease in ERP latencies for both visual and auditory stimuli as infants mature (Chen et al., 2016; Di Lorenzo et al., 2020). These changes likely reflect underlying neurophysiological developments, such as less myelination during infancy which increases with age (Valdés-Hernández et al., 2010).

Future research could benefit from contrasting the mnemonic and inhibitory accounts of infant theta activity. The mnemonic role might be assessed using paradigms that measure looking-time reductions on images as an index of recognition memory, similar to approaches used in non-human primates (Nummela et al., 2019). In contrast, the inhibitory account could be tested by examining the relationship between EEG-measured theta power and fMRI BOLD responses. A negative correlation between theta power and BOLD activity would support the idea that infant theta oscillations reflects functional inhibition(Laufs et al., 2003). An additional promising pursuit would be to investigate dominant frequencies across a broader age span, including children and elderly participants and resting-state activity as well as neural responses to visual stimulation. This would allow us to draw a more complete developmental trajectory of the brain’s oscillatory processing dynamics across the life span.

Compared to previous studies, some adjustments in the paradigm were made to accommodate the infant sample. We presented flickering images of cartoon monsters instead of a flickering light, typically used in adult studies (cf. Herrmann, 2001; VanRullen & Macdonald, 2012). These may have introduced a higher cognitive load. To confirm this approach, we tested adults with the same child-friendly paradigm and replicated the responses in the alpha range for adults. Notably, the higher cognitive load associated with the cartoon stimuli may account for the observed shift in alpha resonance toward 8Hz, rather than the 10Hz frequency observed in previous adult studies (Herrmann, 2001), aligning with previous findings that the alpha rhythm decreases in frequency with higher cognitive load (Klimesch, 1996; Klimesch et al., 1993).

This study is the first to explicitly investigate resonance phenomena in the infant visual system upon rhythmic visual stimulation. We provide convergent evidence of the 4Hz theta rhythm as an inherent rhythm of the infant visual system – unlike adults’ 10Hz alpha rhythm. Our findings suggest that infant brains *tick* at 4Hz, highlighting the importance of the theta rhythm for infant visual perceptual processing and the need to consider neural dynamics when studying the developing brain (Bethlehem et al., 2022; Deen et al., 2017; Ellis et al., 2021; Yates et al., 2025).

Signal to Noise Ratio

## Methods

### Sample

The final sample consisted of 42 8-month-old infants (*MAge* = 7 months, 29 days, *SDAge* = 20 days, *range*[mm;dd]: 07;01 – 09;13, 17 girls) from Regensburg, Germany, born full-term and without record of epilepsy. Eight additional infants were tested but excluded from the final sample due to fussiness (n = 6) or technical errors (n = 2).

Informed written consent was obtained from one parent on behalf of all legal guardians. Participating families received a 10€ voucher for a local toy store. The study was approved by the local ethics committee of the University of Regensburg.

### Stimuli and Procedure

In one laboratory session, infants were presented with the stimuli illustrated in Figure 1 whilst their EEG was recorded. During the experiment, infants sat on their parent’s lap in a darkened room at approximately 70cm from the screen. The stimulus set comprised 75 child-friendly cartoon monsters within a white circle at a visual angle of approximately 14.3 x 14.3°, presented on a gray background. Stimuli were presented at eight discrete frequencies (i.e., appearing and disappearing periodically) or in broadband sequences. For the discrete frequencies, images were flickered sinusoidally at 2, 4, 6, 8, 10, 15, 20 and 30Hz with luminance values between 0 and 100%. This way the average of the luminance of the white circle (switching on and off between black and white) resulted in the same luminance of the gray background, making the stimulation less intrusive to the infants. For the broadband stimulation, sequences with random luminance values were tailored to have equal power at all frequencies. Each infant saw images flickered in 3 distinct broadband sequences (in 4 randomization groups, resulting in a total of 12 different broadband sequences across infants). We presented two additional conditions of either a cartoon monster with no flicker or trials with no image and no flicker (omission condition). Each cartoon monster was presented once in each of the 13 conditions (i.e., eight periodic frequencies, three broadband sequences, no-flicker, omission) for 2 seconds in a random order, continuing until infants lost interest or up to the total of 975 trials in six blocks. Trials were separated by an inter-stimulus interval consisting of a grey screen with a white fixation point for 0.7 to 0.9s. To sustain infants’ attention, each image was paired with one of 20 random ‘monster-sounds’. Each block started with an attention grabber (a cartoon elephant or duck with the corresponding animal sound). On demand, animated videos of rainbows, music notes and trees were manually interspersed. When fussy, some infants were offered with age-appropriate snacks to maintain attention. Stimuli were presented via Psychtoolbox (version 3.0.19.7) and MATLAB (version 2022b). A video recording of the infant was obtained to code and exclude trials that the infant did not watch.

### Electroencephalogram

#### Apparatus

The EEG was recorded continuously with 64 active electrodes (actiCap snap; Easycap GmbH, Herrsching, Germany) positioned according to the international 10-20 system with a target impedance <25kOhm. Two additiona electrodes served as reference (FCz) and ground (FPz). Data were recorded at a sampling rate of 500Hz. Stimuli were presented on a 27-inch screen with a refresh rate of 144Hz.

#### Preprocessing

EEG data were preprocessed and analyzed in MATLAB (Version 2021b) using EEGlab (version 2023.1). In a first step, EEG signals were sampled down to 250Hz, band-pass filtered at 0.5Hz and 45Hz, and segmented into epochs from -1s to 2.5s with respect to image onset. Trials that infants did not attend were identified manually from video recordings and removed. A maximum of 10 noisy EEG sensors were visually identified and interpolated spherically (Köster et al., 2019) and very noisy trials were discarded based on visual inspection. Thereafter, eye-blinks and muscle artifacts were detected using an independent component procedure (ICA; runica algorithm) and removed after visual inspection. Remaining noisy trials were discarded in a second round of visual inspection, and a second ICA was performed to potentially eliminate any remaining eye or muscle artifacts. Finally, the data was re-referenced to the average of all electrodes. In the final sample an average of *M* = 133.5 (*SD* = 59.2) trials remained for the analyses.

### Statistical Analysis

#### Periodic Stimulation

The visual response to periodic stimulation was obtained via Fast Fourier Transformation (FFT) of the signal, computed over the 0 - 2s time window of each trial with a frequency resolution of 0.5Hz. The resulting power spectra were averaged across all trials for a given discrete stimulation frequency. To normalize the spectra, we applied signal-to-noise ratio (SNR) normalization, dividing the power at each frequency by the average power of its two adjacent frequencies (+/- 1Hz; i.e., two frequency steps apart to avoid edge effects; Köster et al., 2019). For example, the SNR at 6 Hz was computed as the power at 6 Hz divided by the average power at 5 Hz and 7 Hz.

Analyses focused on posterior electrodes showing peak RVS response (Oz, Iz, O2, O1, POz, PO4, PO3, PO8, POz). To test harmonic responses, we performed one-sample t-tests against SNR = 1 (chance level). To test whether a significant response emerged in the theta frequency range, we additionally tested the SNR at 4Hz +/- 0.5Hz of each stimulation frequency against SNR = 1. The same analyses were conducted for the no-flicker and omission condition. Note that the omission analysis included only *N* = 41 infants, as one participant did not contribute sufficient trials. To ensure that responses were specific to the theta range, we also tested all frequencies in the alpha range (8-14Hz) against SNR = 1. Finally, to analyze frontal processes, a cluster of frontal electrodes (Fp1, Fz, F3, F7, F4, F8, Fp2, AF7, AF3, AFz, F1, F5, F6, AF8, AF4, F2) was used.

#### Broadband Stimulation

Standardized EEG data from a single trial was cross-correlated with the broadband sequence presented at that particular trial (Chota et al., 2023; VanRullen & Macdonald, 2012). For this purpose, the EEG signal was down-sampled to 144Hz to match the refresh rate of the presentation screen. All time points from the EEG signal between 0 and 2s and the entire duration of the stimulus (2s) were used for calculating the cross-correlation, which was computed at all lags between -2 and 2s. This cross-correlation procedure was aimed at estimating the ‘impulse response function’ (IRF) of the EEG as follows:

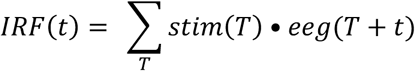

where *stim* denotes the z-scored broadband sequence and *eeg* the z-scored EEG response (van Rullen & Macdonald, 2012; Chota et al., 2023). The results were subsequently averaged over trials.

To evaluate the significance of the IRFs, we generated a Surrogate Response Function (referred to as ‘surrogate’ in the following) as the reference distribution. To this end, we cross-correlated the EEG of a given infant with the broadband sequences presented to the other infants.

Time-frequency representations of the IRFs (and surrogate) were calculated via wavelet transformations. Specifically, we estimated the power spectra by convolving with 7-cycle wide complex Morlet wavelets at the EEG sensors of interest for all frequencies in the 1 to 50Hz range (cf. Chota et al., 2023). The procedure is illustrated for two representative subjects in Figure 3A.

To avoid any condition bias, subject-specific EEG sensors of interest were defined based on mean power across both conditions (IRF and surrogate): For each subject and posterior electrode (CPz, CP2, CP4, CP6, TP8, TP10, Pz, P2, P4, P6, P8, POz, PO4, PO8, Oz, O2, Iz, CP1, CP3, CP5, TP7, TP9, P1, P3, P5, P7, PO3, PO7, O1), we determined the mean cross-correlation across IRF and surrogate and subsequently calculated the spectral power of this condition mean for the entire time window (0 to 2s lag). We then selected the electrode with the greatest 4Hz power of the condition mean and its surrounding five electrodes (in Euclidean distance) for each subject individually. All subsequent analyses of were conducted on these clusters.

The power of the IRF at 4Hz was statistically assessed through comparison to the 4Hz power of the surrogate in the 0 to 1s time lag window with a paired samples t-test (cf.VanRullen & Macdonald, 2012).

To assess whether the IRF reflected reverberations of the full broadband sequence or selectively of the 4Hz component, we conducted a control analysis in which the broadband sequence was notch-filtered around 4Hz. Specifically, we applied a Butterworth band-stop filter (3.5 to 4.5Hz) to remove the 4Hz component while preserving the remaining broadband content. We subsequently recomputed the cross-correlation between the band-stop filtered sequence and the corresponding EEG signal.

#### Adjustments to Analysis of Adult Data

As in the infant sample, we identified EEG sensors of interest for each subject individually based on mean 4Hz power across IRF and surrogate (see above for details). Due to the large interindividual differences in alpha frequencies (Klimesch, 1999), we determined the peak alpha frequency (8-14Hz) for each adult subject, based on the condition mean. That is, we calculated the average SNR across all eight periodic stimulation frequencies and identified the frequency in the 8-14Hz range with the highest average SNR for each subject. Subsequent analyses were conducted with this individual alpha peak frequency.

## Acknowledgments

We would like to thank the infants, their parents and the adult participants for taking part in this study. We are also grateful to Leyla Danabas and Saskia Hermenau for their valuable support with data collection.

## Funding

This study was funded by the Deutsche Forschungsgemeinschaft (DFG; grant number: KO 6028/1-1).

## Author contributions

Conceptualization: MK, MB, OJ Design: MB, MK

Research: MB Analyses: MB, MK, OJ

Writing - original draft: MB, MK

Writing - review and editing: MB, MK, OJ

## Competing interests

Authors declare no competing interests.

## Data and materials availability

Data in accordance with data protection regulations will be made available before acceptance.

## Notes

### Competing Interest Statement

The authors have declared no competing interest.

### Summary of Updates

This version of the manuscript has been revised to include minor edits to the Introduction and Discussion to improve clarity and precision of the language, as well as small adjustments to the framing of the results. No changes were made to the data, analyses, or main conclusions.

