## Supplemental Materials for "Infant Brains Tick at 4Hz – Resonance Properties of the Developing Visual System"

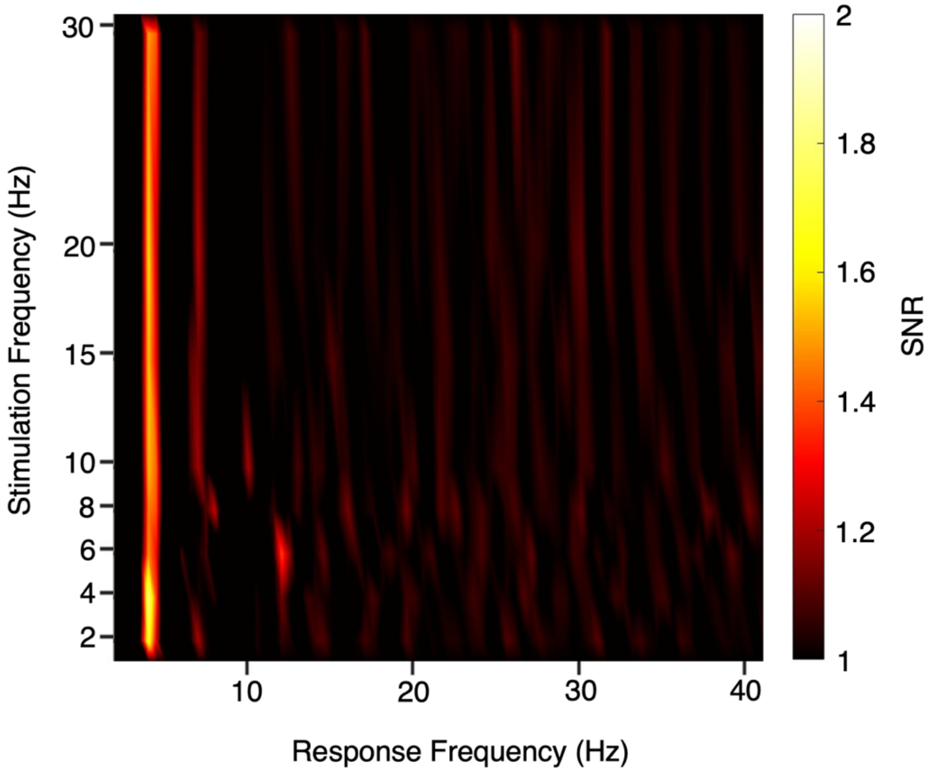


**Supplementary Figure S1. SNR response to periodic visual stimulation at 2, 4, 6, 8, 10, 15, 20, 30Hz at frontal electrodes**. The 4Hz response was particularly pronounced for frontal sites (Fp1, Fz, F3, F7, F4, F8, Fp2, AF7, AF3, AFz, F1, F5, F6, AF8, AF4, F2), where it showed a higher response than first harmonic responses, $t(41) = -3.599$, $p < .001$, and also compared to the frontal 4Hz activity for the no-flicker and omission condition $t(40) = -3.606$, $p < .001$.

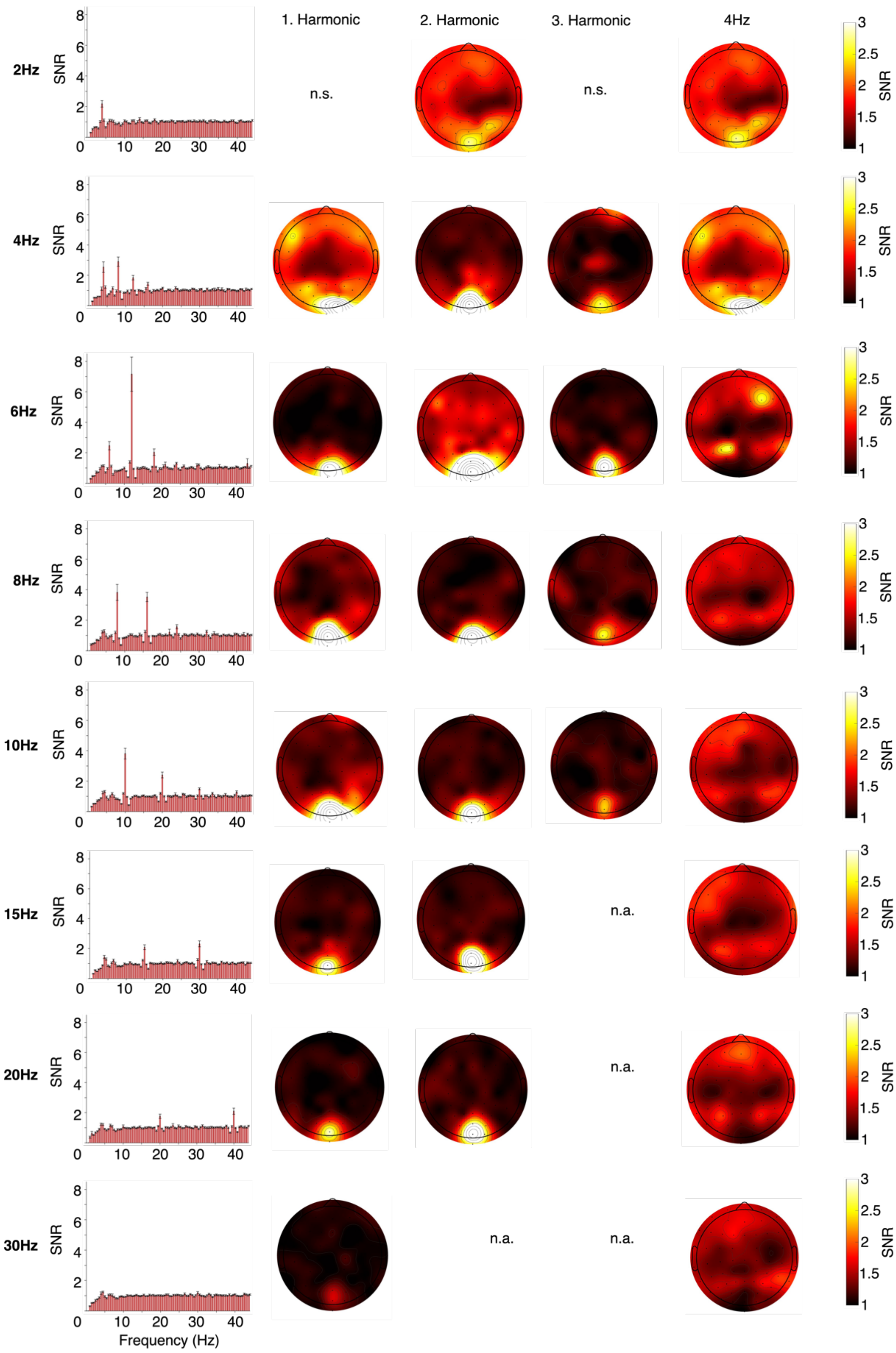


**Supplementary Figure S2. SNR-normalized frequency spectra and topographies for periodic stimulation frequencies (from top to bottom: 2, 4, 6, 8, 10, 15, 20, and 30 Hz).** The left panel shows the SNR-spectrogram for each of the eight discrete stimulation frequencies. The right panel shows the topographic maps of the first, second and third harmonic, and at 4 Hz.

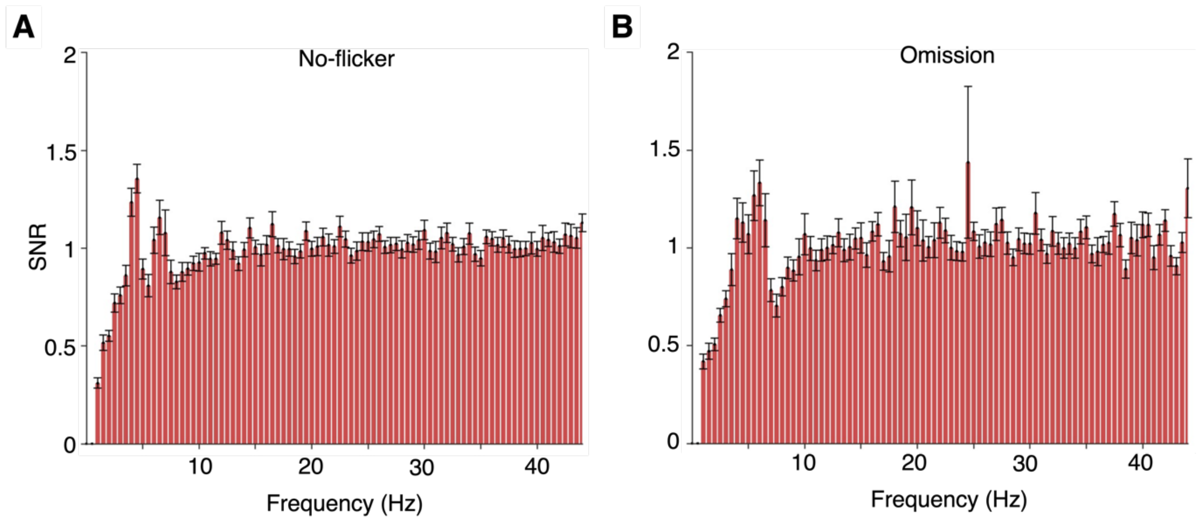


**Supplementary Figure S3. SNR-normalized frequency spectra for the no-flicker and omission condition**. (A) Frequency responses to trials in which an image was presented without flickering. (B) Responses following the presentation of a blank screen. One-sample t-tests against an SNR = 1 revealed significant increases in SNR-corrected power at 4 Hz and 4.5 Hz for the no flicker condition: $t(40) = 3.287$, $p = .001$ and $t(40) = 4.874$, $p < .001$, respectively. In contrast, no significant effects were observed for the omission condition: $t(40) = 1.460$, $p = .08$ (4 Hz) and $t(40) = 1.306$, $p = .10$ (4.5 Hz). *Note:* $N = 41$ participants were included in these analyses, as one infant did not contribute sufficient trials.
